# Products of the Parkinson’s-disease related glyoxalase DJ-1, D-lactate and glycolate, support mitochondrial membrane potential and neuronal survival

**DOI:** 10.1101/006916

**Authors:** Yusuke Toyoda, Cihan Erkut, Francisco Pan-Montojo, Sebastian Boland, Martin P. Stewart, Daniel J. Müller, Wolfgang Wurst, Anthony A. Hyman, Teymuras V. Kurzchalia

## Abstract

Parkinson’s disease is associated with mitochondrial decline in dopaminergic neurons of the *substantia nigra*. One of the genes, DJ-1/PARK7, linked with the onset of Parkinson’s disease, belongs to a novel glyoxalase family and influences mitochondrial activity. It has been assumed that glyoxalases fulfill this task by detoxifying aggressive aldehyde by-products of metabolism. Here we show that supplying either D-lactate or glycolate, products of DJ-1, rescues the requirement for the enzyme in maintenance of mitochondrial potential. We further show that glycolic acid and D-lactic acid can elevate lowered mitochondrial membrane potential caused by silencing PINK-1, another Parkinson’s related gene, as well as by paraquat, an environmental toxin known to be linked with Parkinson’s disease. We propose that DJ-1 and consequently its products are components of a novel pathway that stabilizes mitochondria during cellular stress. We go on to show that survival of cultured mesencephalic dopaminergic neurons, defective in Parkinson’s disease, is enhanced by glycolate and D-lactate. Because glycolic and D-lactic acids occur naturally, they are therefore a potential therapeutic route for treatment or prevention of Parkinson’s disease.

## Introduction

Parkinson’s disease is caused by inexorable deterioration of dopaminergic neurons from the *substantia nigra* (Corti et al., 2011). Although little is known about the onset of Parkinson’s disease, one clue is that a number of genes associated with it are linked to mitochondrial activity (Corti et al., 2011; Federico et al., 2012). One of such genes is DJ-1/PARK7, which was originally reported as an oncogene (Nagakubo et al., 1997), and associated with familial Parkinson’s disease (Bonifati et al., 2003). DJ-1 deficiency was reported to lead to abnormal mitochondrial morphology and dynamics, increased sensitivity to oxidative stress, decreased mitochondrial membrane potential, and opening of the mitochondrial permeability transition pore (Irrcher et al., 2010; Giaime et al., 2012). DJ-1 protein exerts its neuroprotective function against oxidative stress primarily in mitochondria (Junn et al., 2009). Although DJ-1 is predicted and reported to have activity as protease and chaperon (Mizote et al., 1996; Bonifati et al., 2003; Shendelman et al., 2004; Gautier et al., 2012), it is unclear whether these activities contribute to mitochondrial fitness.

DJ-1 was recently reported to belong to a novel glyoxalase family (Lee et al., 2012). Glyoxalases are enzymes that can transform 2-oxoaldehydes glyoxal and methylglyoxal into corresponding 2-hydroxyacids glycolate and D-lactate, respectively. Glyoxal and methylglyoxal covalently react with proteins or lipids to form advanced glycation end-products (AGEs), which are implicated in neurodegenerative diseases including Parkinson’s disease (Castellani et al., 1996; Li et al., 2012). So far, two systems of glyoxalases have been described: 1) Glutathione-dependent Glo I and Glo II systems (Thornalley, 2003) and 2) cofactor-independent Glo III system (DJ-1) (Misra et al., 1995; Lee et al., 2012). Because substrates of glyoxalases are aggressive aldehydes produced by oxidation of glucose during glycolysis (methylglyoxal) and peroxidation of fatty acids (glyoxal), it is assumed that the major function of glyoxalases is to detoxify aldehyde by-products of metabolism (Thornalley, 2003). However, this view has not always been prevalent. Glyoxalases, and their corresponding products (e.g. D-lactate) were considered major components of glycolysis (Ray and Ray, 1998). With the elucidation of the Embden-Meyerhof-Parnas pathway of glycolysis, production of D-lactate was considered an artifact of a biochemical procedure or an undesired side product of glycolysis. Thus, the cellular role of the products of glyoxalases remains unclear.

Here we show that in both HeLa cells and *C.elegans*, the products of DJ-1, glycolate and D-lactate, are required to maintain mitochondrial membrane potential. Remarkably, D-lactate and glycolate increase *in vitro* survival of primary dopaminergic neurons from Parkinson’s model mice embryos. We propose that the products of the glyoxalases are components of a novel pathway that maintain high mitochondrial potential during cellular stress, and that production of glycolate and D-lactate is required to prevent degeneration of dopaminergic neurons in the *substantia nigra*.

### Results

Recently we reported that the *Caenorhabditis elegans* dauer larva, an arrested stage specialized for survival in adverse conditions, is resistant to severe desiccation (Erkut et al., 2011). However, this requires a preconditioning step at a mild desiccative environment (98% relative humidity) to prepare the organism for harsher desiccation conditions (60% relative humidity). We found that during preconditioning, glyoxalase genes *djr-1.2* and *glod-4* were very strongly upregulated ((Erkut et al., 2013); Figure 1A, B, Figure 1-figure supplement 1). We asked whether glyoxalases are required to survive desiccation stress. To address this question, we first produced *djr-1.1;djr-1.2* double mutant missing both DJ-1 homologs (*ΔΔdjr*), and *djr-1.1*;*djr-1.2;glod-4* triple mutant defective additionally in Glo I/II system (*Δglo*) (see Methods). Dauer larvae of these mutants were first preconditioned, and then further desiccated at 60% relative humidity. Subsequently they were rehydrated and measured for their survival rate (Figure 1A, C). The *Δglo* mutant showed significantly increased sensitivity to 60% relative humidity compared to wild type. For instance, 76% of wild type dauers recovered from this stress, while only 22% of *Δglo* did. This result shows that glyoxalases are required for survival after desiccation stress.

**Figure 1.**
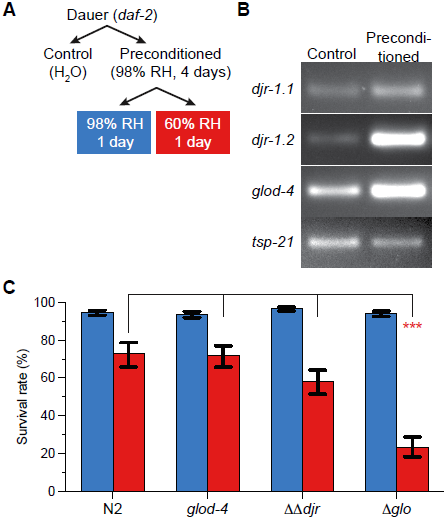
Glyoxalases are required for desiccation tolerance in the *C. elegans* dauer larva. **(A)** Experimental procedure for preconditioning and further desiccation of *C. elegans* dauer larvae. RH, relative humidity. **(B)** Differential expression of *djr-1.1, djr-1.2*, and *glod-4* genes in the *C. elegans* dauer larva upon preconditioning. *Tsp-21* gene was used as the internal control. **(C)** Survival rates of desiccated dauer larvae. Columns show the estimated means of triplicates. Blue, 98% RH; Red, 60% RH. Error bars, standard error of the estimate. *** p < 0.001.

**Figure 1-figure supplement 1.**
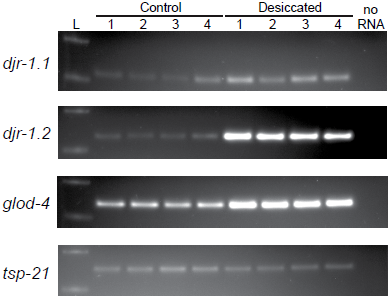
Upregulation of glyoxalase genes upon desiccation of *C. elegans* dauer. The differential expression of *djr-1.1, djr-1.2* and *glod-4* was tested by RT-PCR in four replicates. See Figure 1A for the procedure. *tsp-21* was a control whose expression did not change by desiccation stress.

How does DJ-1 contribute to survival under stress conditions? DJ-1 is reported to exert its neuroprotective function in mitochondria (Junn et al., 2009). Many genes involved in Parkinson’s disease, among them DJ-1, have been linked to alterations in mitochondrial structure and function and an enhanced sensitivity to mitochondrial toxins like Complex-I inhibitors (Clark et al., 2006; Park et al., 2006; Irrcher et al., 2010; Kamp et al., 2010; Sai et al., 2012; Wang et al., 2012; Burchell et al., 2013). Thus we decided to test the structure and function of mitochondria in the absence of DJ-1 (*ΔΔdjr)* or complete glyoxalase activity (*Δglo),* as revealed by staining with MitoTracker CMXRos (Figure 2A) (Pendergrass et al., 2004). In wild-type (N2, top left), mitochondria exhibited elaborated networks, whose intensity of staining represent mitochondrial membrane potential (arrows, Figure 2A) (Pendergrass et al., 2004). Single mutant *glod-4* or *ΔΔdjr* greatly reduced networks. In the triple mutant (*Δglo*) the staining and thus the membrane potential of mitochondria was negligible, and only gut granules (arrowheads) were visible. Thus we conclude that glyoxalases are essential to maintain the mitochondrial structure, potential and function under desiccation stress.

One possibility to explain the data so far presented is that the lack of glyoxalases could lead to the build up of their substrates, toxic aldehydes, leading to phenotypic alteration. However, we propose another hypothesis: the defects may result not only from a build up of toxic aldehydes, but also from the lack of the enzymatic products themselves (α−hydroxyacids). To support this idea, we looked at the effects of the products of glyoxalases, D-lactic acid (DL) and glycolic acid (GA) (Thornalley, 2003; Lee et al., 2012) on the structure and activity of mitochondria. There is no easy way to supply these compounds to dauer larvae, because dauer do not feed. Therefore we decided to use HeLa cells as an alternative model for studying the role of glyoxalases in mitochondrial function.

In HeLa cells, DJ-1 RNAi specifically decreased expression of DJ-1 protein (Figure 2-figure supplement 1) and decreased mitochondrial membrane potential as probed with a J-aggregate forming lipophilic dye JC-1 (Figure 2B) (Smiley et al., 1991), consistent with the previous reports (Larsen et al., 2011; Giaime et al., 2012; Heo et al., 2012). Remarkably, addition of 1 mM GA or DL but not L-lactate (LL) restored mitochondrial membrane potential, while these substances did not affect control Luciferase (Luc) RNAi treated cells (Figure 2B). These results suggest that GA and DL, products of glyoxalases, are required for the activation or maintenance of mitochondrial membrane potential.

**Figure 2.**
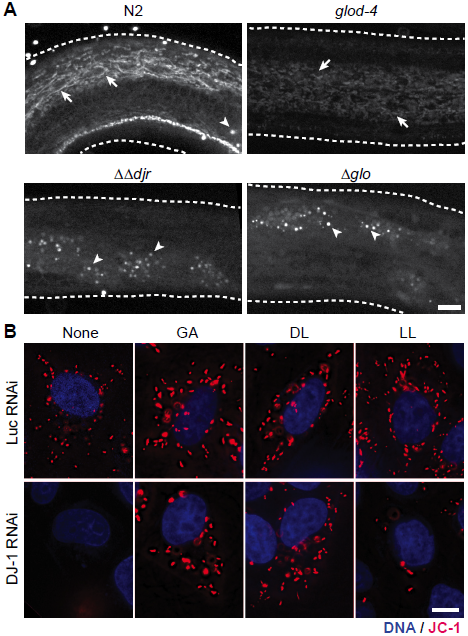
Glycolate and D-lactate are required for mitochondrial membrane potential. **(A)** Disruption of the mitochondrial network upon desiccation at 98% RH and rehydration in mutants for glyoxalases. Arrow, mitochondrial networks stained with MitoTracker. Arrowhead, non-specific staining of the gut granules. **(B)** JC-1 staining in live HeLa cells with 1 mM of glycolate (GA), D-lactate (DL), and L-lactate (LL). Red, JC-1; blue, DNA. For merged images and quantification, see Figure 4. Scale bars, 10 *µ*m.

**Figure 2-figure supplement 1.**
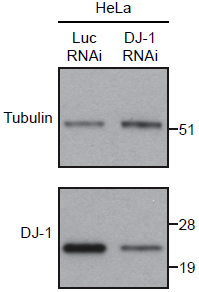
DJ-1 downregulation by esiRNA. Expression of DJ-1 protein in esiRNA-transfected HeLa cells. DJ-1 and tubulin proteins in the whole lysate were detected by immunoblotting. By a densitometric analysis, RNAi of DJ-1 downregulated its expression by 75% in HeLa cells.

Paraquat is an environmental poison known to affect mitochondria (Palmeira et al., 1995); It has been implicated in the onset of Parkinson’s disease, and has been shown to decrease mitochondrial membrane potential (McCarthy et al., 2004; Mitsopoulos and Suntres, 2011). In our assay, paraquat indeed decreased mitochondrial membrane potential in both control and DJ-1 RNAi cells as tested by JC-1 (Figure 3A). Remarkably, addition of GA or DL, but not LL, restored mitochondrial membrane potential of the paraquat-treated cells. In addition to affecting mitochondrial potential, paraquat also induces change in mitochondrial structure (Ueda et al., 1985). Thus we imaged structure of mitochondria in the presence of paraquat with MitoTracker, as it robustly stained mitochondria in HeLa. Although DJ-1 RNAi in HeLa cells did not produce an altered mitochondrial structure (Figure 3-figure supplement 1), together with a low dose of paraquat in DJ-1 RNAi cells, mitochondria became more circular (Figure 3B, insets). This circular phenotype was also rescued by the addition of DL and GA (Figure 3B, insets, Figure 3-figure supplement 1).

**Figure 3.**
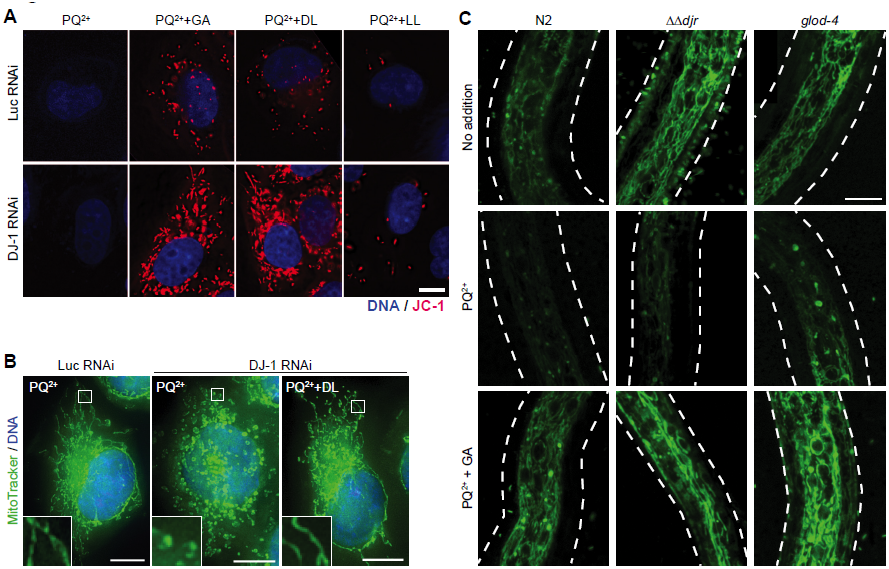
Glycolate and D-lactate rescue the stress-induced mitochondrial defects. **(A)** JC-1 staining in live esiRNA-transfected HeLa cells treated with 50 *µ*M paraquat (PQ^2+^) and 1 mM of glycolate (GA), D-lactate (DL), and L-lactate (LL). Red, JC-1; blue, DNA. For merged images, see Figure 4A. **(B)** Mitochondria of paraquat-treated HeLa cells. Green, MitoTracker; Blue, DNA. Inset, 4x magnification of the boxed area. **(C)** Mitochondria in worm glyoxalase mutant larvae. Wild type (N2), DJ-1 mutant (*ΔΔdjr*) and GLOD-4 mutant (*glod-4*) were treated with PQ^2+^ with or without 10 mM GA, and stained with MitoTracker (green). Dashed lines show the outlines of larvae. Scale bars, 10 *µ*m.

**Figure 3-figure supplement 1.**
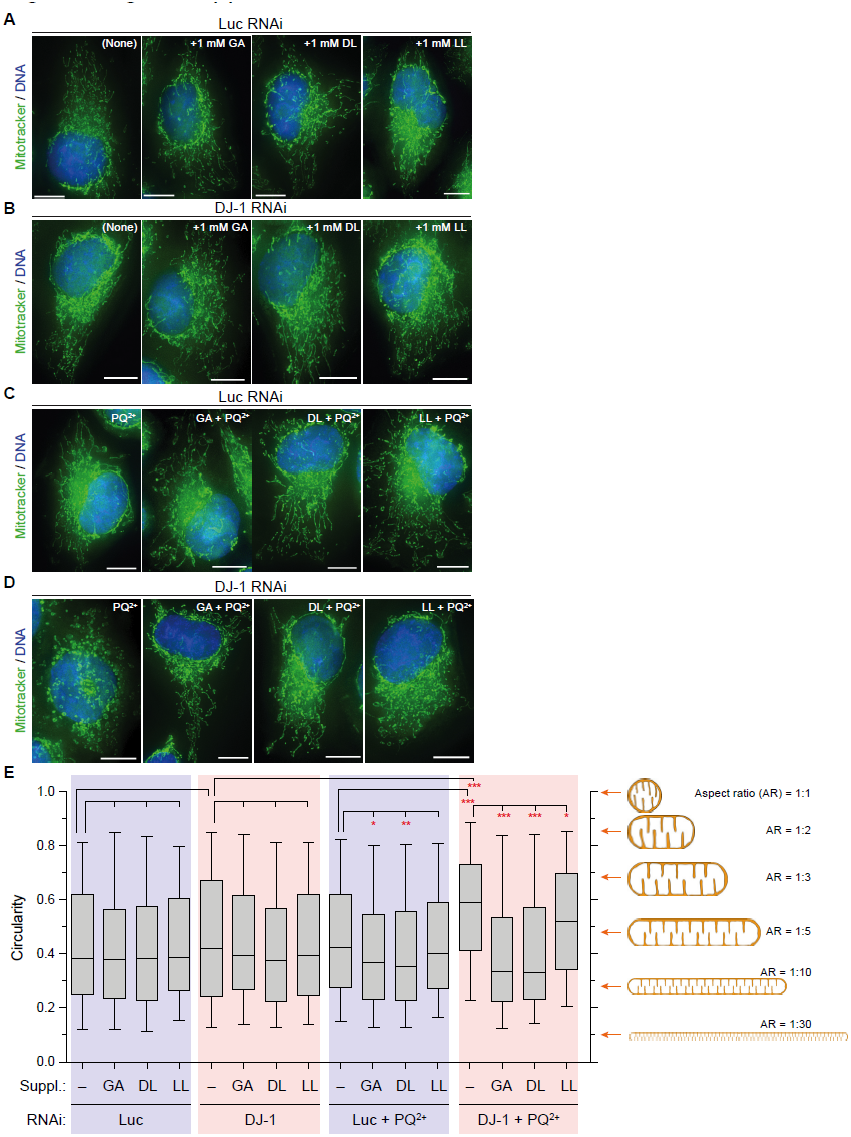
Glycolate and D-lactate rescue mitochondria structure of paraquat-treated DJ-1 RNAi HeLa cells. (A-D) Mitochondria stained with MitoTracker (green) and DNA (blue) of cells treated with control Luciferase RNAi (**A**), DJ-1 RNAi (**B**), control RNAi and paraquat (PQ^2+^) (**C**), and DJ-1 RNAi and PQ^2+^ (**D**). Scale bar, 10 *µ*m. **(E)** Quantification of the mitochondrial network structure. Circularity of mitochondria in cell periphery was calculated (*n* ≥ 280 for each box). On the right, the relation between the mitochondrial shape and circularity is drawn. Circularity in each condition was compared to its own control by one-way ANOVA followed by Tukey’s HSD test. * p < 0.05; ** p < 0.01; *** p < 0.001.

Increasing doses of paraquat resulted in shorter lengths of wild-type and *ΔΔdjr* and *glod-4* mutant reproductive larvae of *C. elegans*, suggesting inhibition of development or growth of worms (Figure 3-figure supplement 2A, B). *ΔΔdjr* mutant exhibited slightly increased sensitivity to paraquat (Figure 3-figure supplement 2C). The decreased viability was rescued by addition of GA (Figure 3-figure supplement 2C). Similarly to human cells, paraquat disrupted mitochondrial membrane potential in reproductive larvae of *C. elegan*s, which was rescued by the addition of GA (Figure 3C).

**Figure 3-figure supplement 2.**
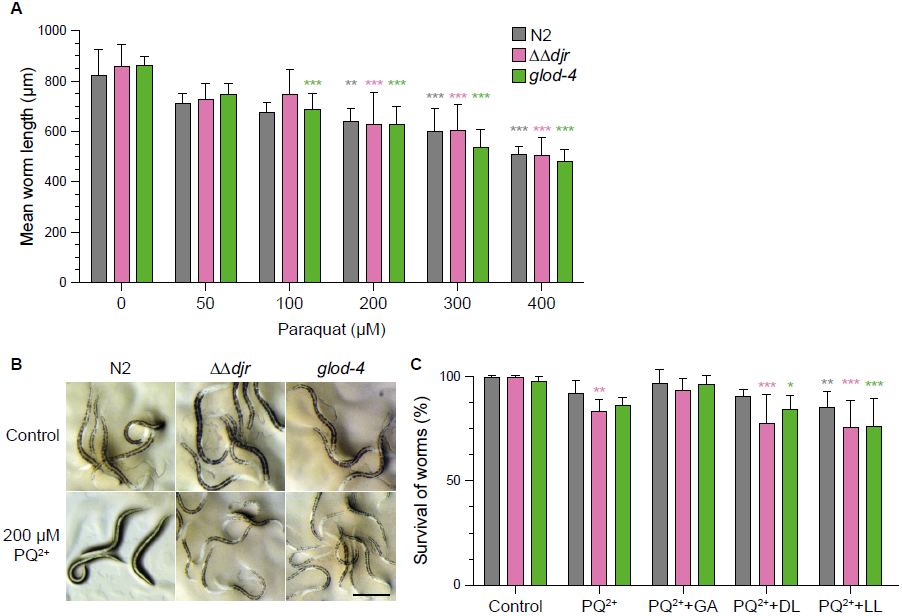
Effects of paraquat on worm larvae. **(A)** Length of the worms treated with paraquat (PQ^2+^). Bars and error bars show the mean and SD, respectively. Sensitivity to PQ^2+^ was comparable between strains (F = 2.334, df = 2, p = 0.1) but overall increased by concentration (F = 81.159, df = 5, p < 0.001). Every strain was compared to its control at different PQ^2+^ concentrations by two-way ANOVA followed by Tukey’s honestly significant differences (HSD) test. **(B)** Worm larvae treated with PQ^2+^ or control. Scale bar, 250 *µ*m. **(C)** Survival of the worm larvae treated with 200 *µ*M paraquat and 1 mM of the indicated supplements. Bars and error bars show the mean and SD, respectively. Every strain was affected differently upon each treatment (strain level F = 10.748, df = 2, p < 0.001; treatment level F = 24.467, df = 5, p < 0.001). PQ^2+^ decreased viability of *ΔΔdjr* mutant, which was restored by glycolate (GA), but not by D-lactate (DL), L-lactate (LL). Viability of *glod-4* was not affected by PQ^2+^ significantly, however the lethality was rescued in a similar way as *ΔΔdjr*. Every strain was compared to its own control by two-way ANOVA followed by Tukey’s HSD test. Data were normalized by Freeman-Tukey’s double arcsine transformation prior to ANOVA. * p < 0.05; ** p < 0.01; *** p < 0.001.

We wanted to test whether GA and DL can rescue a mitochondrial defect caused by loss of other Parkinson’s genes, i.e. PINK1 and Parkin. These genes are genetically and functionally related to DJ-1 (Exner et al., 2007; Hao et al., 2010; Irrcher et al., 2010). Because Parkin was not detected in HeLa (Matsuda et al., 2010), PINK1 was investigated. The protein was downregulated by RNAi (Figure 4-figure supplement 1). PINK1 RNAi decreased mitochondrial membrane potential, as reported previously (Exner et al., 2007). Remarkably, GA and DL rescued this defect (Figure 4A, B). In addition, the substances restored mitochondrial membrane potential of paraquat-treated PINK1 RNAi cells.

**Figure 4.**
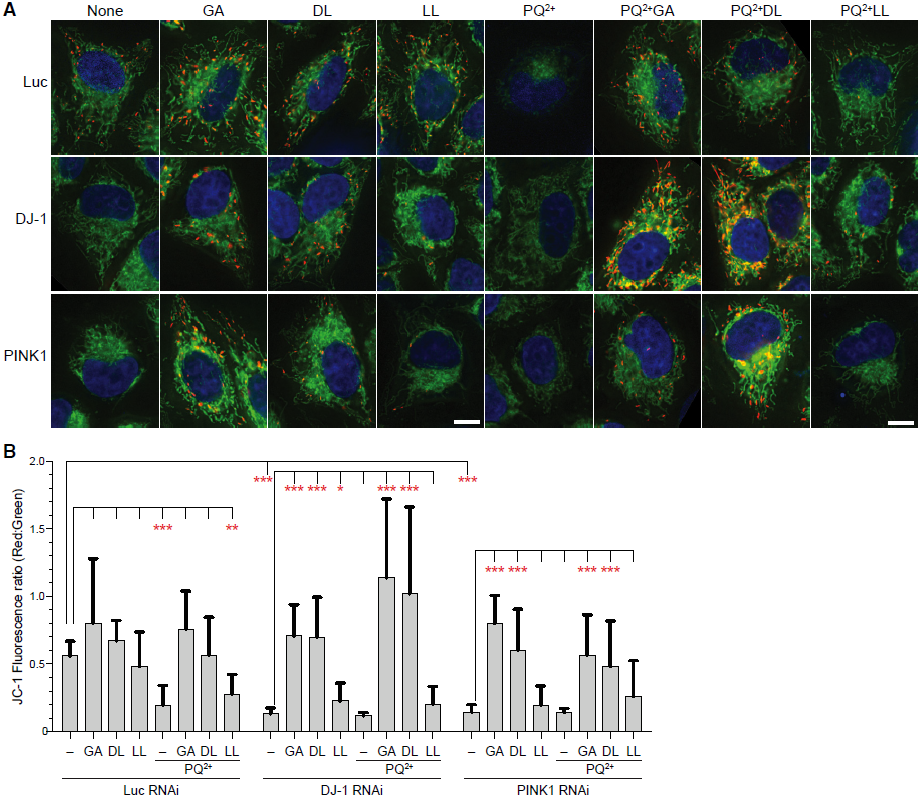
Glycolate and D-lactate rescue lowered mitochondrial membrane potential caused by silencing PINK1. **(A)** Merged images of JC-1 fluorescence in live esiRNA-transfected HeLa cells. Blue, DNA; Green and red, mitochondria with low and high membrane potentials, respectively. Scale bars, 10 *µ*m. (B) The ratio of JC-1 fluorescence intensities in the individual cells. The column and the bar represent the mean and SD, respectively. *n* ≥ 6. * p < 0.05; ** p < 0.01; *** p < 0.001.

**Figure 4-figure supplement 1.**
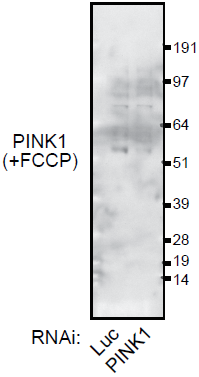
PINK1 downregulation by esiRNA. Expression of PINK1 protein in esiRNA-transfected HeLa cells. Before harvest, cells were incubated with 10 *µ*M FCCP (Carbonyl cyanide 4-(trifluoromethoxy)phenylhydrazone) for 1 hour. Loading of same amounts of total proteins was confirmed by Ponceau S staining (not shown). A 54 kDa band near to the expected molecular weight (63 kDa) was diminished in the PINK1 RNAi sample.

So far, we have shown that the addition of products of glyoxalases can rescue lowered mitochondrial membrane potential caused by downregulation of DJ-1 or PINK1 or by environmental stress. But are GA and DL produced endogenously? We asked whether activation of DJ-1, or glyoxylases in general, during stress leads to a bulk accumulation of DL or GA. For this purpose, we looked for both these compounds in *C. elegans* as well as in HeLa cells. By using a chiral column in an LC-MS application, we could separate D- and L-lactate very effectively (arrows in Figure 4-figure supplement 2A). In dauer larvae before preconditioning, almost all of lactate was present as the L-stereoisomer (Figure 4-figure supplement 2A). Preconditioning did not increase D-lactate to any significant extent. Interestingly, the amount of L-lactate decreased 3-fold, perhaps due to entry of the metabolite into gluconeogenesis. We could not detect any GA in the same amount of extract, while an increased amount of trehalose was detected upon preconditioning, consistent with a previous report (Figure 4-figure supplement 2B, C) (Erkut et al., 2011). Thus we conclude that either these substances are transient intermediates of a pathway needed for the activation of mitochondria, or they are produced in very small amounts and act as signaling molecules.

**Figure 4-figure supplement 2.**
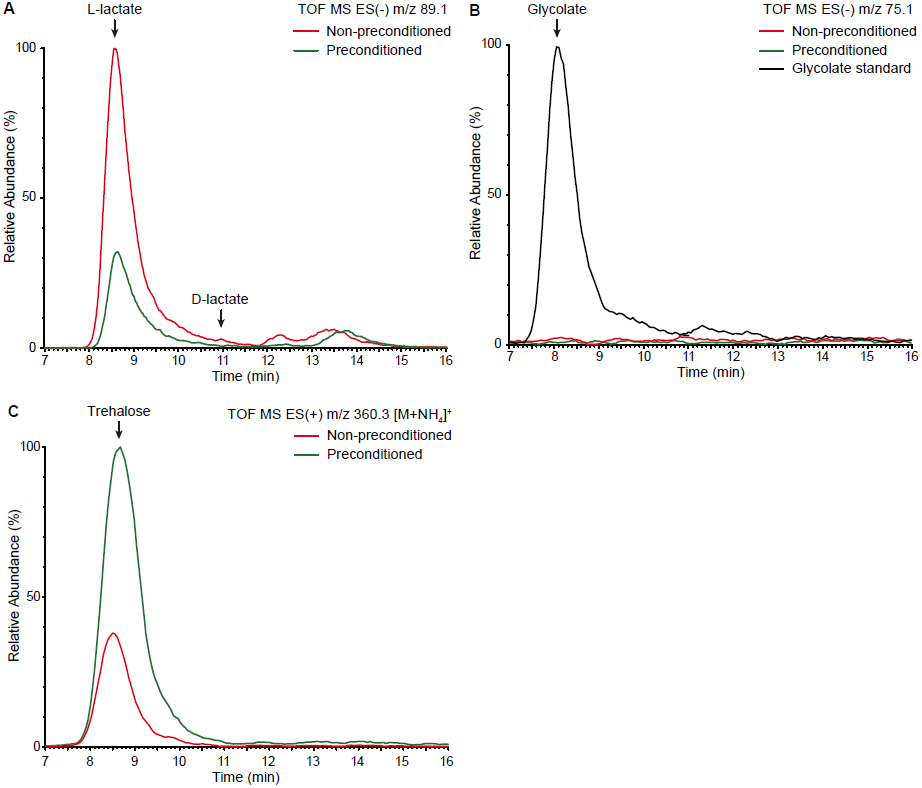
Detection of α-hydroxy acids and trehalose in worms before and after preconditioning. Selected ion monitoring (SIM) chromatograms for molecules of interest are overlaid. **(A)** Separation of lactic acid stereoisoforms. L-lactate decreases 3-fold upon preconditioning whereas D-lactate is in trace amount in worms and its abundance is not affected by desiccation stress. **(B)** No glycolate is detected in worms both before and after preconditioning. **(C)** Trehalose level is increased more than 2-fold upon preconditioning.

We have shown that the products of glyoxalases are required for maintenance of mitochondrial potential in HeLa cells and *C. elegans*. Because toxins that affect mitochondrial function can hasten Parkinson’s disease (Bove et al., 2005), we wondered whether GA and DL would protect neurons against mitochondrial damage. We generated embryonic mesencephalic primary neuron cultures and analyzed the survival of tyrosine hydroxylase-positive (TH^+^) neurons by immunostaining. Strikingly the *in vitro* survival of dopaminergic neurons was stimulated by GA, and DL, but not by LL (Figure 5A, Figure 5-figure supplement 1A). Furthermore, GA and DL significantly rescued the toxic effect of paraquat on the TH^+^ neurons (Figure 5B). We also investigated the effect of these substances on dopaminergic neurons from DJ-1 knock out mice (Figure 5-figure supplement 1B). Similarly, GA and DL could stimulate neuronal survival. These results stress the importance of GA and DL in survival of dopaminergic neurons.

**Figure 5.**
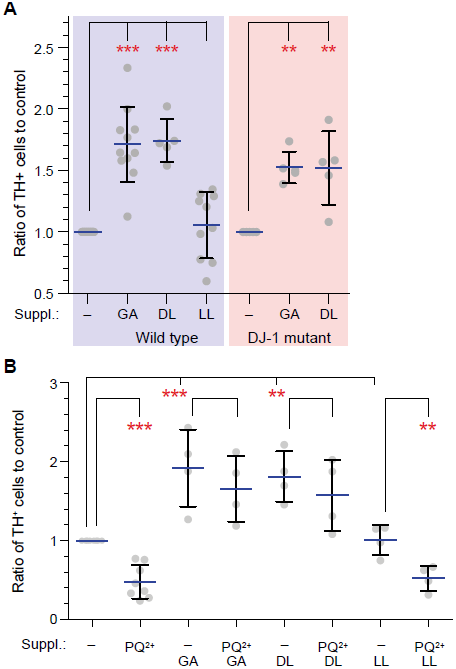
Glycolate and D-lactate support viability of dopaminergic neurons *in vitro*. **(A)** Viability of the dopaminergic neurons *in vitro* from wild type and DJ-1 mutant mice embryos in the presence or absence of 10 mM glycolate (GA), D-lactate (DL), or L-lactate (LL). Points show the ratio of tyrosine hydroxylase-positive (TH^+^) cells to the control (no supplement). Horizontal blue bars and error bars represent the mean and SD, respectively. ** p < 0.01; *** p < 0.001. **(B)** Viability of the dopaminergic neurons in the presence of 12.5 *µ*M paraquat (PQ^2+^) in the absence or presence of 10 mM GA, DL, or LL. Primary neurons isolated from wild-type E14.5 embryos were cultured *in vitro* in the presence of the indicated supplements and PQ^2+^. Points show the ratio of TH^+^ cells to the control. Horizontal blue bars and error bars represent the mean and SD, respectively.

**Figure 5-figure supplement 1.**
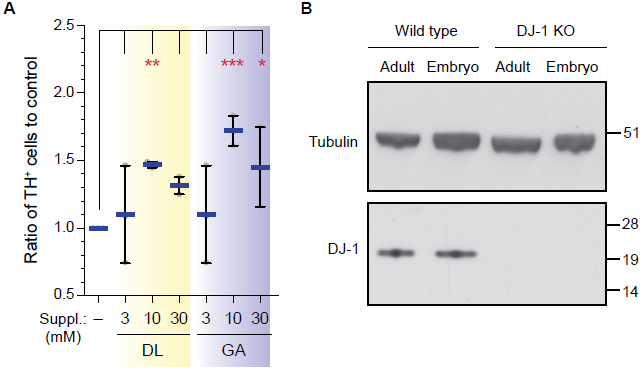
Glycolate and D-lactate support *in vitro* survival of the dopaminergic neuron. **(A)** Survival of the primary dopaminergic neurons in the presence of the different concentrations of D-lactate (DL) and glycolate (GA). The primary neurons from wild type mouse embryos were cultured with the indicated substances for 6 days, fixed, and stained for tyrosine hydroxylase (TH), a dopaminergic neuron-specific marker. The relative number of the TH^+^ cells to none-treated control was plotted, with mean (blue) and SD. Each dot indicates an independent experiment. **(B)** Immunoblot of DJ-1 in wild type and DJ-1 mutant mouse brains. Total brains isolated from wild type and DJ-1 mutant adults and embryos were lysed, and tested for DJ-1 expression. Tubulin is the loading control.

## Discussion

In this paper, we have shown that the products of glyoxalases, glycolate and D-lactate, are required to maintain mitochondrial membrane potential. Maintenance of mitochondrial membrane potential is associated with response to desiccation stress in worms, and importantly, the survival of dopaminergic neurons in mammals. Our data therefore highlight an understudied aspect of the Embden-Meyerhof glycolytic pathway: A small fraction of triose-phosphate is converted into methylglyoxal, which is further transformed into D-lactate by glyoxalases (Thornalley, 2003). It has so far been thought that glyoxalases protect cells by removing products of glycolysis or lipid oxidation. Thus, our data allow us to suggest the idea that glyoxalases have two functions: on one hand they detoxify chemically aggressive aldehydes and on the other hand they produce compounds necessary for maintaining mitochondrial potential.

DJ-1 is a member of a new class of glutathione-independent glyoxalases that have recently been identified, and studies of their enzymology are just beginning. *In vitro* enzyme assays using NMR have shown that both human and *C.elegans* DJ-1 can produce glycolic acid and lactate, but they did not distinguish between L and D lactate (Lee et al., 2012). More recently, studies on the biochemical activity of DJ-1 expressed *in vitro*, using enzyme-coupled assays, suggest that it has only a weak activity as a methylglyoxalase (Hasim et al., 2014). Indeed, our measurements of D-lactate and glycolate (see Figure 4-figure supplement 2) show that they do not accumulate in high amounts, which further suggests that they do not function in high concentration. However, it is also possible that these glyoxalases use other substrates more efficiently. Further studies will be required to understand the *in vivo* regulation and activity of the DJ-1 family, but this will require improved methods to detect flux through the glyoxalase pathway *in vivo*. However the facts that D-lactate and glycolic acid have such an effect on mitochondria, and rescue the DJ-1 phenotype, strongly suggest that these substances are important products of the DJ-1 glyoxalase *in vivo*.

We do not yet understand how D-lactate and glycolate increase or maintain mitochondrial potential. For instance it is quite possible that these products are further processed. We know, however, that GA and DL restore mitochondrial membrane potential in cells depleted of DJ-1, PINK1 or treated with paraquat. Thus, GA and DL are core compounds in a general pathway that maintains mitochondrial potential. Because both paraquat and loss of DJ-1 are thought to increase permeability of the inner mitochondrial membrane (Costantini et al., 1995; Giaime et al., 2012), one possibility is that GA and DL decrease permeability of the mitochondrial membrane under stressed conditions. Understanding this molecular mechanism is an avenue for future investigation.

It seems likely that the symptoms of Parkinson’s disease, neuronal cell death in the *substantia nigra*, arise from an increased sensitivity of dopaminergic neurons to diminished mitochondrial membrane potential. (Corti et al., 2011; Federico et al., 2012). A decline in mitochondrial activity would therefore tend to exacerbate this problem. Indeed, recent experiments in *C. elegans* show that mitochondria are required for survival of neurons (Rawson et al., 2014). Further investigation of the phenotypes we observe should clarify which aspects of cellular metabolism require mitochondrial potential. Our data show that there is no reduction in ATP levels in DJ-1 mutant conditions, suggesting that it is not an energy requirement (data not shown). Rather, other mitochondrial metabolism cycles, such as one-carbon metabolism (Tibbetts and Appling, 2010; Locasale, 2013) may be involved. Other pathways, such as the glyoxylate shunt in worms, may also play a role.

Therapeutic routes for Parkinson’s disease have so far been symptomatic and intractable. It has been shown that environmental toxins that affect mitochondria are strongly linked to the appearance of Parkinson’s disease (Song et al., 2004; Freire and Koifman, 2012) and impairment of the mitochondrial function is a common feature of both idiopathic and genetic Parkinson’s disease (Schapira et al., 1989; Clark et al., 2006; Park et al., 2006; Irrcher et al., 2010; Kamp et al., 2010; Pan-Montojo et al., 2012; Wang et al., 2012; Braidy et al., 2013; Burchell et al., 2013). Our discovery that the production of molecules from endogenous enzymatic pathways can protect neurons, offers a potential therapeutic direction that could include preventive strategies. Both products of glyoxalases exist in many natural products. Thus, providing neurons with these substances might protect them against metabolic or environmental stress. Because many diseases are associated with a decline in mitochondrial activity (Schapira, 2012), the products of glyoxalases could have a general role in protecting cells from decline.

## Materials and Methods

### Chemicals

Glucose (Merck), glycolic acid (15451, ACROS Organics) neutralized with NaOH to pH = 7.4, and sodium D-lactate (71716, Sigma), glyoxal (128465, Sigma), paraquat (sc-257968, SantaCruz biotechnologies or 36541 Fluka^®^ from Sigma-Aldrich) were used.

### Cell culture, RNAi, worm strains

HeLa cells were maintained in DMEM (Life Technologies) supplemented with 10% fetal bovine serum (FBS), 2 mM GlutaMAX, 100 unit/ml penicillin, 100 **µ**g/ml streptomycin at 37°C in a 5% CO_2_ environment. HeLa cells were RNAi-transfected with 10 nM endoribonuclease-digested small interfering RNA (MISSION esiRNA, Sigma) using Oligofectamine RNAi MAX reagent (Invitrogen). Cells were assayed after 48 hours from RNAi.

All *C. elegans* strains were maintained on NGM agar plates seeded with *Escherichia coli* NA22 at 15°C (Brenner, 1974). Mutant strains *djr-1.1*(*tm918*), *djr-1.2*(*tm951*) and *glod-4*(*tm1266*) were obtained from National Bioresource Project, Japan. Wild type (N2) and *daf-2*(*e1370*) mutant strains were obtained from *Caenorhabditis* Genetics Center, USA. All mutants were outcrossed at least twice with the wild type to eliminate background mutations.

### Preparation, culture and treatment of primary mesencephalic dopaminergic neurons from mouse embryos

Primary mesencephalic neuronal cell cultures were prepared as previously described (Gille et al., 2004). Briefly, brain mesencephalons from E14.5 C57JBL6 or DJ-1 embryos were dissected under the microscope and digested with Trypsin-EDTA (Sigma-Aldrich). The trypsin reaction was stopped by adding the basic medium (BM) containing Neurobasal A medium (Gibco), 1 mg/ml penicillin/streptomycin, 10% (v/v) fetal calf serum (Invitrogen) and 2 mM L-Glutamine and cells were mechanically dissociated using a fire-polished Pasteur pipette. Medium was fully replaced by centrifuging for 5 min. at 1200 rpm, aspiring the supernatant and adding 8 ml of the fresh BM to the pellet. Concentration of cells in the medium was estimated and cells were plated in a volume of 250 *µ*l in 4-well plates (176740, Nunc, Thermo Scientific) or 35 *µ*l in *µ*-clear 96-well plates (Greiner) coated with poly-D-lysine (Sigma-Aldrich) at a concentration of 2 × 10^6^ cells per ml. The same volume of medium containing the different treatment substances was added 4 hours after plating to obtain the following treatment concentrations: control, 3 mM glucose, 10 mM GA, 10 mM DL and 10 mM LL. 24 hours later 1/3 of the medium was replaced with fresh BM. On DIV3 (day-in-vitro-culture 3) half of the medium was replaced with B27 medium containing Neurobasal A medium, 1 mg/ml penicillin/streptomycin, 2 mM L-Glutamine (Sigma-Aldrich) and B-27 supplement (Life Technologies) and on DIV5 all medium was replaced by B27 medium. On DIV7 cell were either fixed using Accustain^®^ (Sigma-Aldrich) for 30 min. or PQ^2+^ treated at a concentration of 12,5 **µ**M for 72 hours more and fixed.

### Immunocytology of mesencephalic cell cultures

Accustain^®^ fixed neuronal cell cultures were washed 3 × 10 min. in phosphate buffered saline (PBS), blocked using a blocking solution (BS) (0.2% Triton X-100 in PBS and 5% donkey serum (DS)) for 1 hour at RT, and incubated with mouse anti-TH (1:500, Millipore), chicken anti-ßIII-tubulin (1:500, Millipore) and rabbit anti-TOM20 (1:200, FL-145, Santa Cruz Biotechnology) or rabbit anti-NeuN (1:500, Millipore) primary antibodies in BS overnight at 4 °C. On the next day cells were washed 4 × 10 min. with PBS, incubated in donkey Alexa^®^ 488 anti-rabbit, donkey Alexa^®^ 555 anti-mouse (Life Technologies) and donkey Alexa^®^ 647-anti-chicken (Jackson Immunoresearch) secondary antibodies for 1 hour at RT, washed 4 × 10 min. with PBS, incubated with Hoechst33342 for 10 min. and washed once more in PBS.

### Generation of *C. elegans* multiple mutant strains

*djr-1.1* and *djr-1.2* males and hermaphrodites were first crossed reciprocally. L4 hermaphrodites from the F_1_ generation were singled out and let lay eggs for 2 days. Subsequently, the adults were lysed and genotyped individually.

One adult was put in 100 *µ*l lysis buffer (1X PCR buffer and 200 ng/*µ*l proteinase-K in water), snap-frozen in liquid nitrogen and incubated for 1 hour at 65 °C. Then, the enzyme was denatured at 98 °C for 15 min. Genotyping PCR was performed in 1X PCR buffer with MgCl_2_, 200 *µ*M dNTP mix, 400 nM of each primer, 0.02 U *Taq* polymerase and 5 *µ*l of gDNA from the lysis of an adult hermaphrodite using the primers listed in Supplementary file 1. PCR conditions were the following: Initial denaturation at 94 °C for 10 min, amplification in 30 cycles of 94 °C for 30 sec, 62 °C for 25 sec and 72 °C for 30 sec, final extension at 72 °C for 10 min.

Populations arising from an individual heterozygous for both alleles were selected and L4 hermaphrodites were singled out for one more round of genotyping as described above. Finally, 3 lines homozygous for both alleles were found. One of these lines was selected to be used in subsequent experiments. We named this double mutant *ΔΔdjr*.

**Supplementary file 1.**
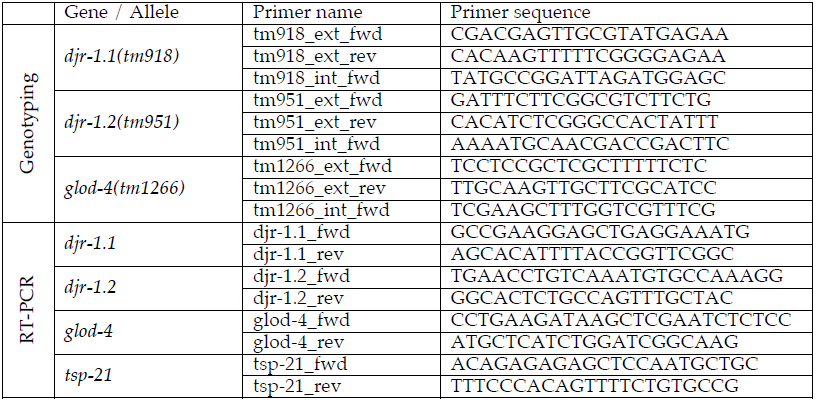
Sequences of the primers used in genotyping and RTPCR.

As a next step, *ΔΔdjr* mutants were crossed with *glod-4(tm1266)* mutants. The genotyping and selection of homozygous triple mutants were done similarly, using the primers listed in Supplementary file 1. PCR conditions were the same as above, except that the annealing temperature was increased to 65 °C. Finally, two lines homozygous for all 3 alleles were obtained. One of these lines was used in experiments. For convenience, we named this triple mutant *Δglo*.

### Desiccation and measurement of desiccation tolerance in dauers

N2, *glod-4, ΔΔdjr* and *Δglo* dauers were generated on agarose plates by sterol depletion/lophenol substitution (Matyash et al., 2004). *daf-2* dauers were generated in liquid culture by growing at 25 °C. Worms were preconditioned at 98% RH for 4 days and/or desiccated at 60% RH for 1 day in controlled humidity chambers (Erkut et al., 2013). Survival rate of different strains were calculated as the percentage of survivors in the total population after rehydration.

### Measurement of desiccation-induced gene expression

Total RNA was extracted from *daf-2* dauers before and after preconditioning in 4 biological replicates. 500 ng of total RNA was reverse transcribed using 250 ng oligo(dT)_12-18_ primer (Invitrogen, CA), 5 nmol dNTP and 200 U SuperScript III reverse transcriptase (Invitrogen, CA) at 50 °C for 1 h according to the protocol supplied by the manufacturer. PCR was performed in 1X PCR buffer with MgCl_2_, 200 *µ*M dNTP, 500 nM of forward and reverse primers, and 0.05 U *Taq* polymerase using 0.1 *µ*l of the cDNA product. PCR conditions were the following: 94 °C for 10 min, 30 cycles of 94 °C for 20 sec, 61.5 °C for 25 and 72 °C for 15 sec, finally 72 °C for 5 min. The primer sequences are presented in Supplementary file 1. PCR products are separated on a 3% agarose gel and visualized using ethidium bromide.

### Paraquat treatment of HeLa cells and worm larvae

For experiments with paraquat (PQ^2+^) in cells, 50 *µ*M PQ^2+^ and 1 mM GA/DL/LL were added to HeLa cells and incubated for 24 hours. The media and the supplements were replaced 1 hour before fixation or assay.

Worms were treated with 200 *µ*M PQ with or without 10 mM GA, and compared to worms that are not treated with either PQ or GA. Subsequently, they were stained and imaged for mitochondrial organization and activity.

### Mitochondrial live staining of worm larva

Mitochondria staining was performed as previously described (Yang and Hekimi, 2010). Briefly, Mitotracker Deep Red and CMXROS (M22426, M7512, Life Technologies) were dissolved in DMSO at a concentration of 5 mM and kept at -20 C as a stock solution. On the day of microscopy worms were incubated in a 1:1000 diluted Mitotracker for 45 min. at room temperature.

Worms were then paralyzed with 1 mM Levamizol (Sigma-Aldrich), placed on slides covered with a thin layer of NGM medium on top of which the coverslip (22 × 22 mm, Menzel-Glaser #1) was fixed using nail lack.

### Light microscopy, and image analysis

To image mitochondria of HeLa cells, MitoTracker Red CMXRos was added at 150 nM and fixed with 3% (v/w) paraformaldehyde in PBS, 1 mM MgCl_2_, and 5 mM EGTA. DNA was counter-stained by 1 *µ*g/ml Hoechst33342. As MitoTracker Red CMXRos robustly stained mitochondria in HeLa, JC-1 (Santa Cruz Biotechnology, 10 *µ*g/ml) was used to visualize mitochondrial membrane potentials in live cells, with 100 ng/ml Hoechst33342. They were imaged on the DeltaVision system using a 60x objective (PlanApo N, NA = 1.42, Olympus) equipped with a CO_2_ supply, and deconvolved and maximally projected images were used for the analysis. Due to high background of MitoTracker in the center of the cell, mitochondria in a 15 × 15 *µ*m area in the periphery were manually annotated on ImageJ software. Total integrated intensity of green- and red–fluorescent JC-1 in the individual cells was measured to obtain the fluorescence ratio. All JC-1 images were adjusted for the contrast in the same way on ImageJ.

Microscopy images from live paralyzed worm larvae stained with Mitotracker were taken using a confocal microscope (LSM510, Zeiss, Germany). Samples were excited using a 514 (Mitotracker CMXRos) or a 647 (Mitotracker Deep Red) nm lasers and two channels, one BP505-550 or LP650 and the BF channel were used to acquire the images. Gain was maintained between 450 and 515 for all samples to ensure the detection of signal intensity differences.

### Counting of dopaminergic neurons

Dopaminergic TH^+^ neurons were observed using an inverted fluorescence microscope (Axiovert 200M, Zeiss) under a 10x objective (PlanApo, NA = 0.45). The diameter of every well was scanned in two perpendicular directions (i.e. top to bottom and left to right) and total TH^+^ neurons were counted for every well.

### Immunoblotting

EsiRNAi-transfected HeLa and mouse brains were lysed in a lysis buffer (50 mM HEPES pH= 7.5, 150 mM KCl, 1 mM MgCl_2_, 10% glycerol, 0.1% NP-40) with a protease inhibitor cocktail (Complete, Roche), resolved in SDS-PAGE, and transferred onto a nitrocellulose membrane. In immunoblotting, the following primary antibodies were used: human DJ-1 (FL-189, SantaCruz, 1:200 dilution); mouse DJ-1 (HPA004190, Sigma, 1:250); PINK1 (38CT20.8.5, Thermo, 1:500); alpha-tubulin (DM1A, Sigma, 1:2000). Horseradish peroxidase-conjugated anti-IgG antibodies (Bio-rad, 1:2000) were used for the secondary antibody. Chemiluminescence by ECL reagent was developed on a Hyperfilm (GE healthcare).

### Liquid chromatography mass spectrometry (LC-MS) analysis of alpha-hydroxy acids

*daf-2* dauers directly collected from the liquid culture or preconditioned at 98% RH for 4 days were homogenized and extracted according to Bligh and Dyer’s method (Bligh and Dyer, 1959). The aqueous phases were dried and dissolved again in 50% methanol (v/v) using volumes calculated according to total soluble protein amounts. HeLa cells harvested directly or treated with GA were also extracted by the same method. The final volume of their aqueous fractions was normalized according to the number of cells.

Separation and detection of glycolic and lactic acids were performed by normal-phase LC-MS using an Agilent G1312A pump equipped with an Agilent Autosampler G1329A. Separation employed a Cogent Diamond Hydride column (25 cm × 4.6 mm i.d., 4 *µ*m, Microsolv) coupled on-line to a Waters/Micromass LCT time of flight (TOF) mass spectrometer equipped with electrospray ionization (ESI). Alpha-hydroxy acid species were separated by isocratic elution. An aqueous formic acid (0.1%; v/v), including 10 mM ammonium formate/acetonitrile mixture (30:70; v/v) was used. The column was operated at 40 °C and flow rate was set to 1 ml/min with a split to 100 *µ*l/min into the mass spectrometer. 5 *µ*l of each sample were injected into the column.

For the separation of D- and L-lactic acid enantiomers, we employed chiral chromatography by using an Astec CHIROBIOTIC R chiral column (25 cm × 4.6 mm i.d., 5 *µ*m, Supelco), as reported previously (Henry et al., 2012). Isocratic elution was performed with the mobile phase 15% (v/v) 30 mM ammonium acetate in H_2_O (adjusted with acetic acid to pH 3) and 85% (v/v) acetonitrile. The column was operated at 5 °C with a split to 50 *µ*l/min into the same mass spectrometer.

The mass spectrometer was operated with a spray voltage of 2.5 kV and a source temperature of 140 °C in negative and positive ion mode. Nitrogen was used as the cone and nebulizing gas at flow rates of approx. 40 and 500 L/h, respectively. Positive and negative ion full scan mass spectra were acquired from the m/z range of 60–1000 mass units in a scan time of 1 s. The system was operated and the resulting data were processed by MassLynx (Version 4.1) software (Waters).

### Statistics and graph representation

Statistical differences between the tested treatments were determined by ANOVA followed by the Tukey’s honestly significant differences post-hoc test. Survival rates were compared using beta regression (Erkut et al., 2013). Data expressed in percentages were first transformed by Tukey’s double arcsine function (Freeman and Tukey, 1950) to achieve normal distribution prior to ANOVA. Statistical analysis and graphs were done on a Prism software version 5 (GraphPad Inc.) and an R environment.

### Acknowledgments

We thank Suzanne Eaton and Marino Zerial at the Max Planck Institute for Molecular Cell Biology and Genetics, for critical reading of the manuscript and constructive suggestions. We thank Frank Buchholz and Mirko Theis for esiRNA, Zoltan Maliga, John Asara and Min Yuan for helpful discussions, Marc Bickle for suggestions on mitochondria annotation, Ina Poser for technical help, and Human Protein Atlas for the DJ-1 antibody.

### Author Contributions

Y.T., C.E., F.P-M., D.J.M., A.A.H. and T.V.K. conceived and designed the experiments. Y.T., C.E., F.P-M., S.B. and M.P.S. performed the experiments. Y.T., C.E., F.P-M., W.W., A.A.H. and T.V.K. interpreted and analyzed the data. Y.T., C.E., F.P-M., A.A.H. and T.V.K. wrote the paper.

